# Imprinted anti-hemagglutinin and anti-neuraminidase antibody responses after childhood infections of A(H1N1) and A(H1N1)pdm09 influenza viruses

**DOI:** 10.1101/2023.02.01.526741

**Authors:** Pavithra Daulagala, Brian R. Mann, Kathy Leung, Eric H.Y. Lau, Louise Yung, Ruipeng Lei, Sarea I.N. Nizami, Joseph T. Wu, Susan S. Chiu, Rodney S. Daniels, Nicholas Wu, David Wentworth, Malik Peiris, Hui-Ling Yen

**Affiliations:** School of Public Health, Li Ka Shing Faculty of Medicine, The University of Hong Kong, Hong Kong SAR, China; WHO Collaborating Center for Surveillance, Epidemiology and Control of Influenza, Centers for Disease Control and Prevention, USA; WHO Collaborating Centre for Infectious Disease Epidemiology and Control, School of Public Health, Li Ka Shing Faculty of Medicine, The University of Hong Kong, Hong Kong SAR, China; Laboratory of Data Discovery for Health Limited (D24H), Hong Kong Science Park, Hong Kong SAR, China; The University of Hong Kong – Shenzhen Hospital, Shenzhen, China; Department of Biochemistry, the University of Illinois at Urbana-Champaign, Illinois, USA; Department of Paediatrics and Adolescent Medicine, Queen Mary Hospital and Li Ka Shing Faculty of Medicine, The University of Hong Kong; The Francis Crick Institute, The Crick Worldwide Influenza Centre, WHO Collaborating Centre for Reference and Research on Influenza, Midland Road, London, UK; Centre for Immunology and Infection (C2I), Hong Kong Science Park, Hong Kong SAR, China

**Author notes:** Correspondence: Malik Peiris and Hui-Ling Yen.

**Keywords:** influenza, imprinting, neuraminidase, hemagglutinin, antigenic drift

## Abstract

Immune imprinting is a driver known to shape the anti-hemagglutinin (HA) antibody landscape of individuals born within the same birth cohort. With the HA and neuraminidase (NA) proteins evolving at different rates under immune selection pressures, anti-HA and anti-NA antibody responses since childhood influenza infections have not been evaluated in parallel at the individual level. This is partly due to the limited knowledge of changes in NA antigenicity, as seasonal influenza vaccines have focused on generating neutralising anti-HA antibodies against HA antigenic variants. Here we systematically characterised the NA antigenic variants of seasonal A(H1N1) viruses from 1977 to 1991 and completed the antigenic profile of N1 NAs from 1977 to 2015. We identified that NA proteins of A/USSR/90/77, A/Singapore/06/86, and A/Texas/36/91 were antigenically distinct and mapped N386K as a key determinant of the NA antigenic change from A/USSR/90/77 to A/Singapore/06/86. With comprehensive panels of HA and NA antigenic variants of A(H1N1) and A(H1N1)pdm09 viruses, we determined hemagglutinin inhibition (HI) and neuraminidase inhibition (NI) antibodies from 130 subjects born between 1950-2015. Age-dependent imprinting was observed for both anti-HA and anti-NA antibodies, with the peak HI and NI titers predominantly detected from subjects at 4-12 years old during the year of initial virus isolation, except the age-independent anti-HA antibody response against A(H1N1)pdm09 viruses. More participants possessed antibodies that reacted to multiple antigenically distinct NA proteins than those with antibodies that reacted to multiple antigenically distinct HA proteins. Our results support the need to include NA proteins in seasonal influenza vaccine preparations.

**IMPORTANCE:** Seasonal influenza vaccines have aimed to generate neutralizing anti-HA antibodies for protection since licensure. More recently, anti-NA antibodies have been established as an additional correlate of protection. While HA and NA antigenic changes occurred discordantly, the anti-HA and anti-NA antibody profiles have rarely been analysed in parallel at the individual level, due to the limited knowledge on NA antigenic changes. By characterizing NA antigenic changes of A(H1N1) viruses, we determined the anti-HA and anti-NA antibody landscape against antigenically distinct A(H1N1) and A(H1N1)pdm09 viruses using sera of 130 subjects born between 1950-2015. We observed age-dependent imprinting of both anti-HA and anti-NA antibodies against strains circulated during the first decade of life. 67.7% (88/130) and 90% (117/130) of participants developed cross-reactive antibodies to multiple HA and NA antigens at titers ≥1:40. With slower NA antigenic changes and cross-reactive anti-NA antibody responses, including NA protein in influenza vaccine preparation may enhance vaccine efficacy. (150 words)

## INTRODUCTION

Influenza viruses continue to pose significant morbidity, mortality, and socioeconomic burden worldwide as antigenic variants that evade pre-existing immunity continuously emerge, causing regular epidemics and infrequent pandemics. The most commonly administered inactivated influenza vaccines induce neutralising antibodies targeting the receptor binding domain of the main surface glycoprotein, hemagglutinin (HA), and this strategy is supported by the establishment of the HA inhibition (HI) antibodies as a key correlate of protection (1). Other serological correlates of protection, such as HA-stalk reactive antibodies (2, 3) and anti-neuraminidase (NA) antibodies (4, 5), have been identified more recently through observational and human challenge studies. Specifically, anti-NA antibodies have been reported to protect against infection, reduce symptoms, and/or shorten the duration of viral shedding (2-7).

Individual and population immunity are continuously shaped by influenza antigenic drifts and shifts. During primary exposure to the influenza virus, memory B cells are developed to react to a range of conserved and non-conserved viral epitopes. These memory B cells may bias future antibody responses upon exposure to an antigenic variant in a phenomenon referred to as “immune imprinting”. Immune response to a subsequent infection or vaccination is thus influenced by the antigenic similarity of later strains to an individual’s initial exposure. Experimental and observational studies have identified immune imprinting as an important driver of the anti-HA antibody immune landscape with both beneficial and disadvantageous impacts (8-11).

The protective effect of immune imprinting has been observed in mice sequentially immunised with antigenic drift variants of A(H1N1) viruses, A/Puerto Rico/8/1934 (PR8) and its antigenic variant A/Puerto Rico/8/1934-S12a (S12a), which resulted in higher affinity antibodies to the primed PR8 HA protein. However, these mice still developed antibodies that reacted to S12a, albeit at a lower HI titer (12). Despite developing a biased antibody response towards PR8, passive transfer of PR8-S12a immune sera protected naïve mice from S12a challenge. On the other hand, epidemiological and modelling studies suggest that immune imprinting provides limited protection against antigenically more distinct variants than the imprinted strain (13, 14) due to structural similarity and conserved protective epitopes unique to either Group 1 HA or Group 2 HA proteins.

The immune imprinting patterns of anti-NA antibodies are relatively less studied due in part to the limited knowledge on the NA antigenic changes over time. Seasonal influenza vaccines have focused on generating neutralising anti-HA antibodies against HA drift variants since the initial licensure in 1945. An observational study reported the age-dependent immune imprinting of anti-NA antibodies using historical N1 and N2 strains that were distantly related in antigenicity (15). It is unknown if age-dependent immune imprinting can be similarly observed using well-defined drift NA proteins as the virus continues to evolve over time. As HA and NA drift variants evolve discordantly (16-18), the age-dependent anti-HA and anti-NA antibody responses have not been systematically compared in parallel. Here, we characterised the NA antigenicity of seasonal A(H1N1) viruses from 1977-1991 to complete the NA antigenic profile of A(H1N1) and A(H1N1)pdm09 viruses circulating in humans. By comparing imprinting patterns of anti-HA and anti-NA antibody responses, a broader cross-reactivity of anti-NA antibody responses than anti-HA antibody responses was observed.

## RESULTS

### Antigenic changes in the NA of A(H1N1) seasonal influenza virus, 1977-1991

To complete N1 NA antigenic mapping since its re-emergence in 1977, we evaluated the NA antigenic changes among A(H1N1) WHO vaccine strains from 1977 to 1991, including A/USSR/90/77 (USSR/77), A/Brazil/11/78 (Brazil/78), A/Chile/1/83 (Chile/83), A/Singapore/6/86 (Singapore/86), and A/Texas/36/91 (Texas/91). The NA inhibition (NI) titers of polyclonal ferret anti-sera raised against the A(H1N1) viruses from 1977 to 1991 were measured against recombinant H6N1 viruses with NA genes derived from the A(H1N1) vaccine strains (19) using the Enzyme-linked Lectin Assay (ELLA) (20). Since the NA coding sequence of Brazil/78 was identical to that of USSR/77, it was not included in the characterisation. A bi-directional change ≥4-fold in NI antibody titers between two strains in two-way HIs was considered antigenically distinct.

According to the two-way analysis (Fig. 1 and Table 1), the ferret anti-sera raised against USSR/77 showed an NI titer of 1:320 against the homologous antigen (Fig. 1a). The ferret anti-sera raised against Chile/83 showed a two-fold reduction in NI titer against NA of USSR/77, while the NI reactivity of ferret anti-sera against USSR/77 showed a four-fold reduction in NI titer against Chile/83 (Fig. 1b). As such, the antigenic change of NA between Chile/83 and USSR/77 was only evident in one direction. The ferret anti-sera raised against Singapore/86 showed a homologous NI titer against its own NA at a dilution of 1:320 (Fig. 1c). The anti-sera for Singapore/86 showed a ≥16-fold and ≥4-fold reduction in NI titer against NA of USSR/77 and Chile/83, respectively. Similarly, the ferret anti-sera raised against older strains (USSR/77 and Chile/83) also showed a ≥4-fold reduction in NI titer against NA of Singapore/86, suggesting an NA antigenic drift from Chile/83 to Singapore/86. The ferret anti-sera raised against Texas/91 poorly inhibited NAs of older strains with a ≥32-fold NI titer reduction, and the ferret anti-sera raised against older strains showed a ≥16-fold NI titer reduction against NA of Texas/91 (Fig. 1d), showing a pronounced change in NA antigenicity from Singapore/86 to Texas/91.

**Table 1.**
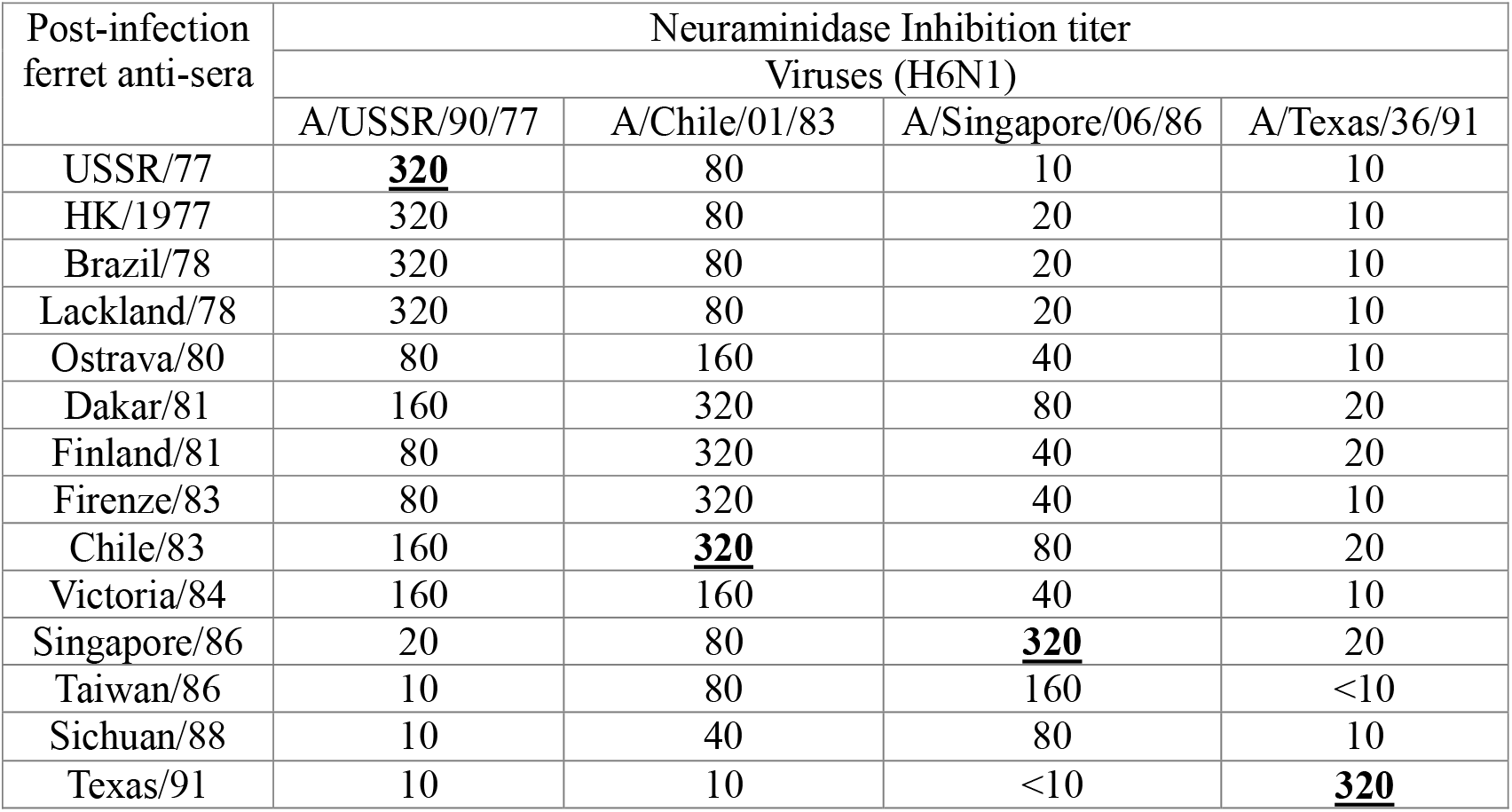
Extended NI analysis with ferret anti-sera raised against additional A(H1N1) viruses.

| Post-infection<br>ferret anti-sera | Neuraminidase Inhibition titer |  |  |  |
| --- | --- | --- | --- | --- |
|  | Viruses (H6N1) |  |  |  |
|  | A/USSR/90/77 | A/Chile/01/83 | A/Singapore/06/86 | A/Texas/36/91 |
| USSR/77 | **<u>320</u>** | 80 | 10 | 10 |
| HK/1977 | 320 | 80 | 20 | 10 |
| Brazil/78 | 320 | 80 | 20 | 10 |
| Lackland/78 | 320 | 80 | 20 | 10 |
| Ostrava/80 | 80 | 160 | 40 | 10 |
| Dakar/81 | 160 | 320 | 80 | 20 |
| Finland/81 | 80 | 320 | 40 | 20 |
| Firenze/83 | 80 | 320 | 40 | 10 |
| Chile/83 | 160 | **<u>320</u>** | 80 | 20 |
| Victoria/84 | 160 | 160 | 40 | 10 |
| Singapore/86 | 20 | 80 | **<u>320</u>** | 20 |
| Taiwan/86 | 10 | 80 | 160 | <10 |
| Sichuan/88 | 10 | 40 | 80 | 10 |
| Texas/91 | 10 | 10 | <10 | **<u>320</u>** |

**Figure 1.**
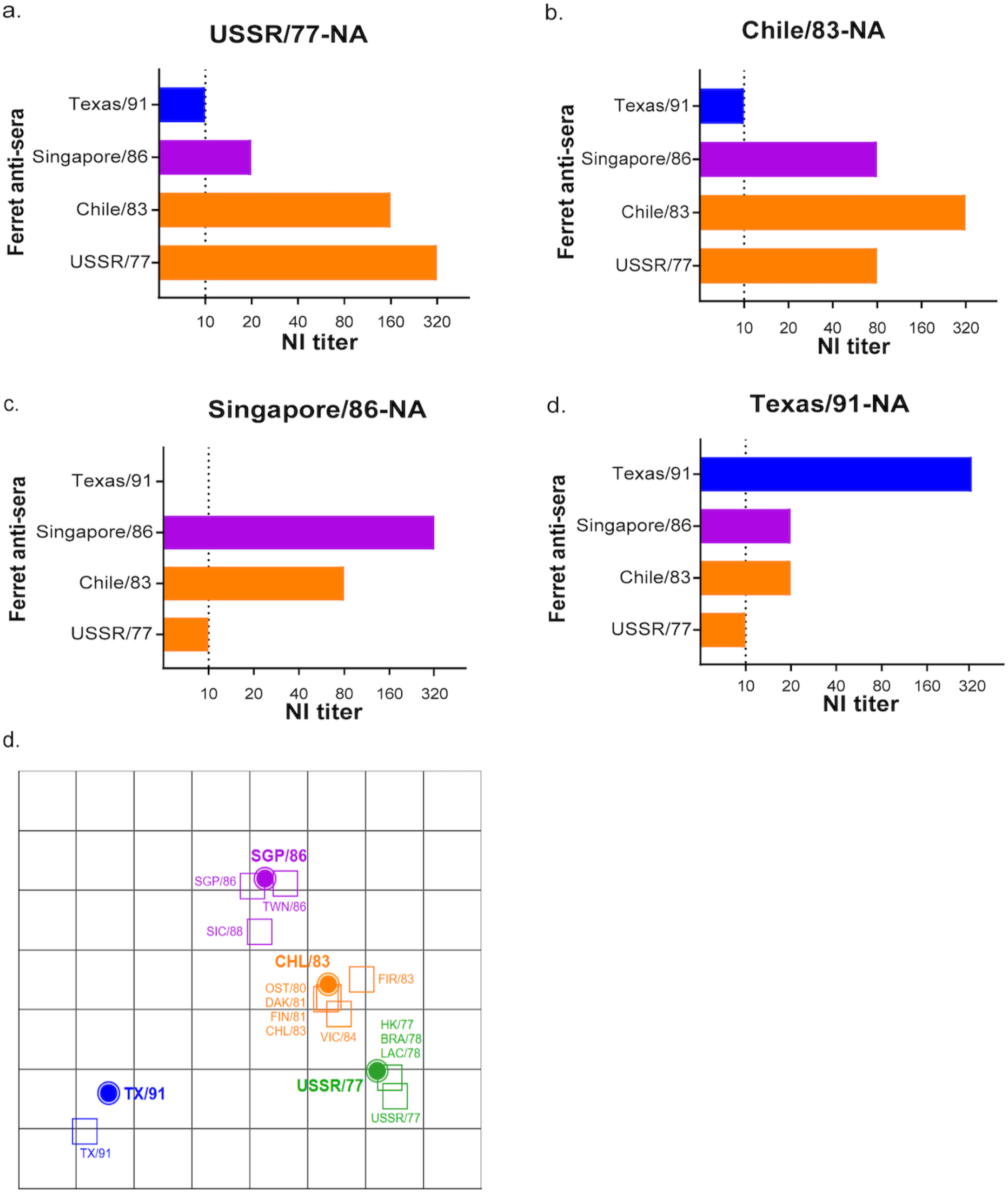
Identification of NA antigenic variants from A(H1N1) viruses circulating between 1977-1991. NI titers were determined using ferret anti-sera and recombinant H6N1 viruses carrying NA protein derived from **a**, USSR/77, **b**, Chile/83, **c**, Singapore/86, or **d**, Texas/91. **e**, Antigenic cartography map was generated using the end-point NI titers as shown in Table 1. NI titer values of 5 were assigned for all results below the ELLA limit-of-detection (NI titer <10). Recombinant A(H6N1) viruses (filled circle icons) and H1N1-raised ferret antisera (open square icons) are color-coded by N1 genetic similarity to USSR/77 (green), Chile/83 (CHL/83; orange), Singapore/86 (SGP/86; purple), and Texas/91 (TX/91; blue). Grid increments between two icons indicate a 2-fold difference in modelled NI titers with two units corresponding to a 4-fold reduction, three units to an 8-fold reduction, *et cetera*.

The antigenicity of N1 protein derived from USSR/77, Chile/83, Singapore/86 and Texas/91 was further validated using available ferret sera raised against other A(H1N1) viruses that circulated in 1977-1988 (Table 1). Anti-sera raised against A(H1N1) that circulated between 1977 to 1984 showed a 4-fold change in NI titers against the NA derived from USSR/77 or Chile/83 viruses but reacted poorly (≥4-fold change) against the NA of Singapore/86 (Table 1). With the available ferret anti-sera, our result showed that A/Victoria/4/84 (Victoria/84) was the latest strain that showed similar antigenicity with USSR/77. Anti-sera raised against Singapore/86 and A/Taiwan/1/86 showed comparable NI titers against different NA proteins, suggesting they share similar antigenicity. Anti-sera raised against A/Sichuan/4/88 showed a four-fold change in NI titer against the NA of Singapore/86 and a 32-fold change in NI titer against the NA of Texas/91, suggesting that its NA protein may be antigenically different from these viruses. Antigenic cartography map (Fig. 1e) produced from the endpoint NI titers of A(H1N1)-raised ferret anti-sera supports the antigenic relatedness of USSR/77 and Chile/83 NA proteins with a one-way antigenic change. In contrast, the NA of Singapore/86 is more antigenically distant to USSR/77 and Chile/83. From this antigenic map, it can be deduced that the NAs of USSR/77, Chile/83 and Singapore/86 evolved sequentially, while the NA of Texas/91 was both antigenically distinct and distant.

### The N386K amino acid change has led to a one-way NA antigenic change from USSR/77 to Singapore/86

Victoria/84 shared antigenicity with USSR/77 but was antigenically distinct from Singapore/86. We hypothesised that a key amino acid change may have occurred during the evolution from Victoria/84 to Singapore/86, leading to antigenic drift. By comparing the NA coding sequences of Victoria/84 and Singapore/86, a total of 15 amino acid changes (including 9 in the NA head domain) were identified (Table 2). The effects of S247N, K369R, N386K and K434N (N1 numbering) amino acid substitutions were further investigated (Fig. 2a, 2b) as they were localised at antigenic sites previously identified for N1 or N2 proteins (18, 21-23).

**Table 2.**
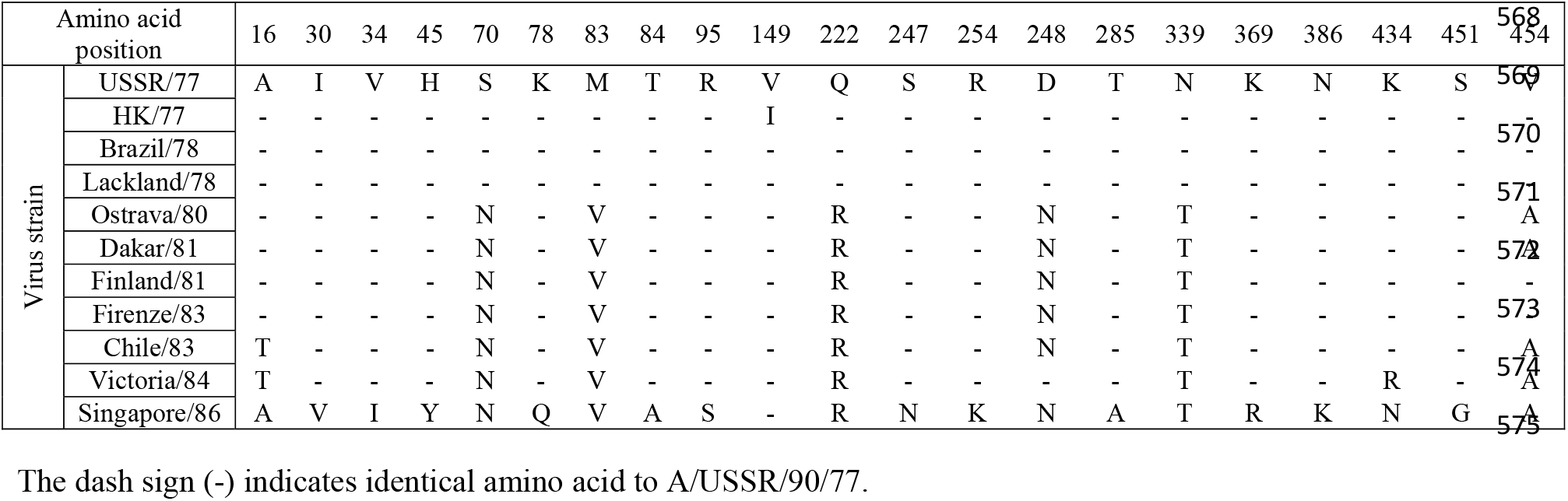
NA Amino acid changes from A/USSR/90/77 to A/Singapore.

**Figure 2.**
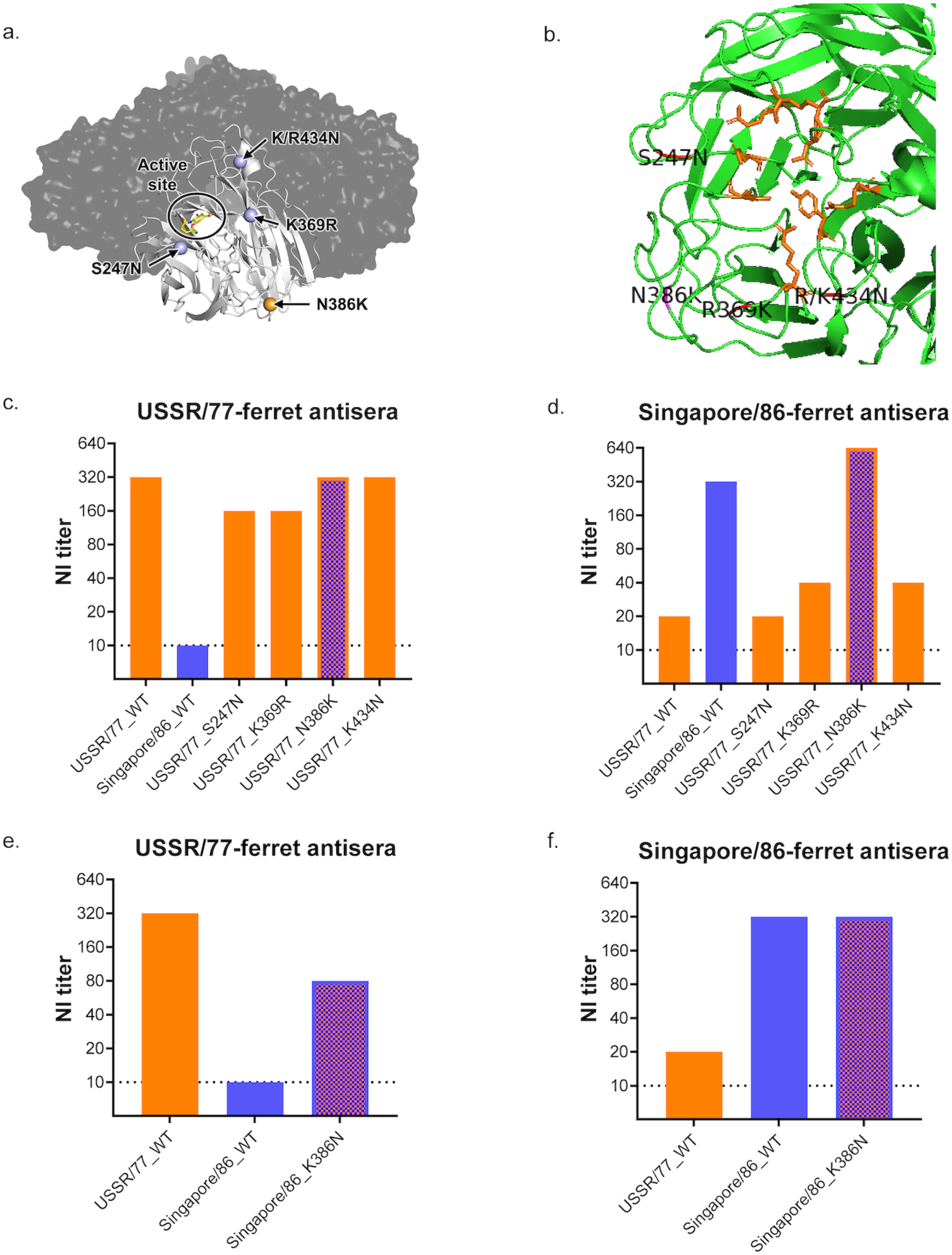
Mapping amino acid changes responsible for the antigenic drift from USSR/77 to Singapore/86. **a**, The NA enzyme active site and key amino acid changes, S247N, K369R, N386K, K/R434N were mapped on the N1 tetramer of A/Vietnam/1203/2004 (PDB 2HU0) using PyMOL. **b**, Magnified view of N1 monomer of A/Vietnam/1203/2004 (PDB 2HU0) with the active site residues highlighted in orange and critical amino acid residues highlighted in pink. **c**, Reactivity of anti-USSR/77 ferret sera against NA protein derived from USSR/77, Singapore/86, or USSR/77 with different amino acid changes. **d**, Reactivity of anti-Singapore/86 ferret sera against NA protein derived from USSR/77, Singapore/86, or USSR/77 with different amino acid changes. **e**, Reactivity of anti-USSR/77 ferret sera against NA protein derived from USSR/77, Singapore/86, or Singapore/86 with K386N substitution. **f**, Reactivity of anti-Singapore/86 ferret sera against NA protein derived from USSR/77, Singapore/86, or Singapore/86 with K386N substitution.

Using site-directed mutagenesis, S247N, K369R, N386K, or K434N amino acid changes were each introduced into the NA gene of the USSR/77 and recombinant A(H6N1) viruses were generated to determine NI titers. Anti-USSR/77 ferret sera reacted similarly to the wild-type (WT) and mutant USSR/77 NA proteins, with NI titers ranged from 1:160 to 1:320, suggesting that the S247N, K369R, N386K or K434N amino acid changes did not significantly affect the binding of anti-USSR/77 ferret sera (Fig. 2c). The WT and mutant USSR/77 viruses were further tested using anti-Singapore/86 ferret sera, which did not react well with the WT or mutant USSR/77 NA proteins carrying the S247N, K369R, or K434N amino acid changes with NI titers ranged from 1:20 to 1:40 (Fig. 2d). However, the Singapore/86 antisera showed comparable NI titers (1:320 to 1:640) against the homologous Singapore/86 NA protein and the USSR/77 protein containing the N368K amino acid change. These results suggest that the change at residue 386 from N (USSR/77) to K (Singapore/86) may have affected the NA antigenicity. We further validated the role of residue 386 by introducing the K386N amino acid change into the NA protein of Singapore/86. Anti-USSR/77 ferret sera reacted well with the USSR/77 NA (NI titer at 1:320) and poorly with Singapore/86 NA (1:10), but the antisera showed increased reactivity against Singapore/86 NA containing the K386N change (1:80) (Fig. 2e). Interestingly, the anti-Singapore/86 ferret sera reacted similarly to Singapore/86 NA proteins with or without the K386N change (Fig. 2f). Taken together, these results showed that the amino acid change at residue 386 was associated with one-way NI titer change.

An NA phylogenetic tree was constructed using representative A(H1N1) viruses circulated from 1977 to 2008, and amino acid changes at NA residue 386 were monitored over time (Fig. 3). A(H1N1) circulating from 1977 to 1984 contained asparagine at NA residue 386, corresponding to the related NA antigenicity of USSR/77 and Chile/83 viruses during this period. The N386K amino acid substitution emerged in 1986, leading to the antigenic drift observed from USSR/77 to Singapore/86. The N386K amino acid substitution was maintained among A(H1N1) viruses isolated from 1986 to 1988. Aspartic acid replaced lysine at NA residue 386 in 1988, prior to the emergence of the antigenically distinct Texas/91. The K386D substitution was observed in majority of the A(H1N1) viruses circulating from 1989 to 2008.

**Figure 3.**
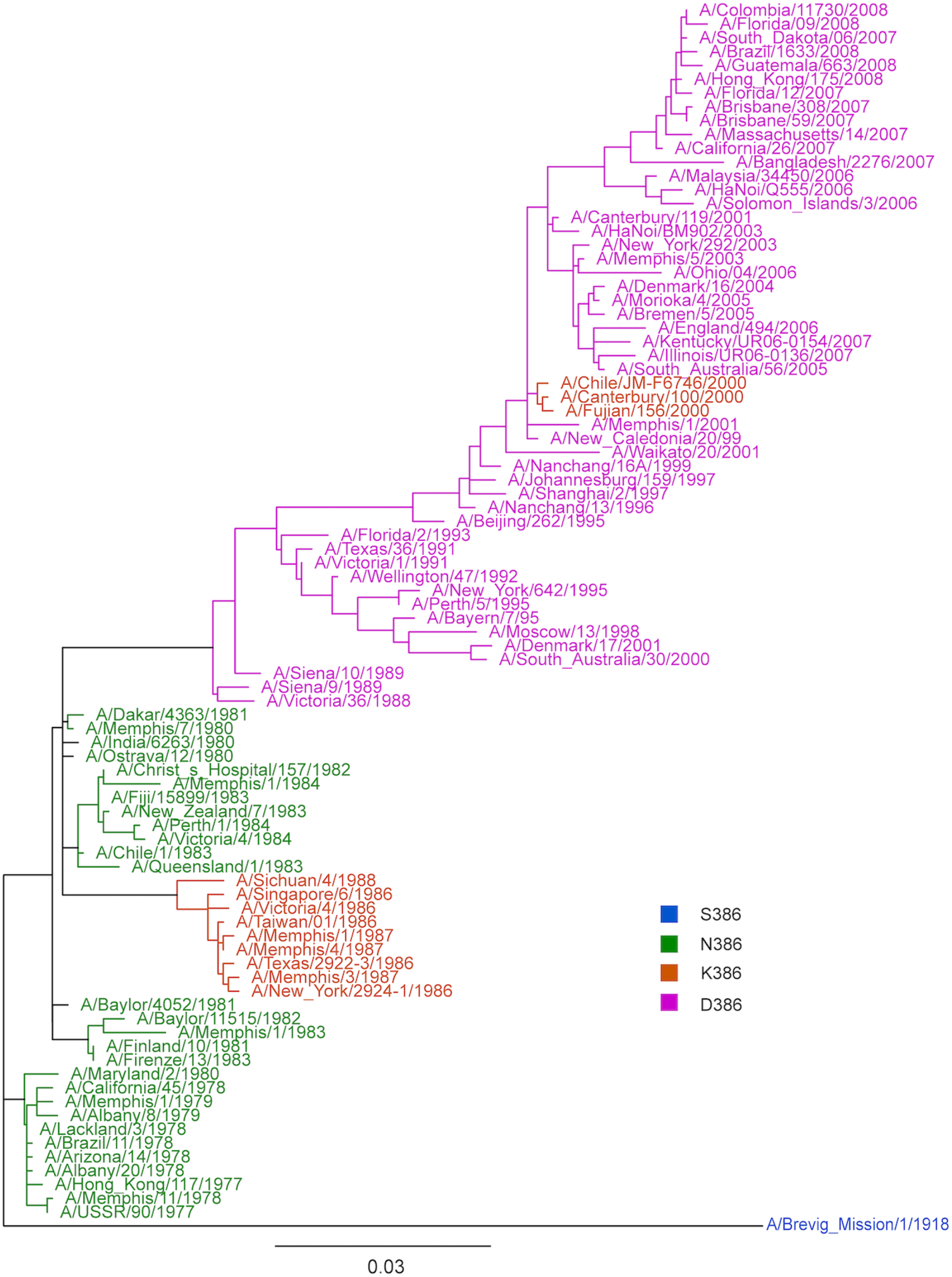
Phylogenetic tree of N1 sequences from seasonal A(H1N1) circulated from 1977-2008. A phylogenetic tree was generated using the full coding regions of the NA proteins from representative strains from 1977 to 2008. The strains are colour-coded depending on the amino acid substitution on residue 386. The viruses with amino acid substitution of serine (S), asparagine (N), lysine (K) and aspartic acid (D) at residue 386 are shown in blue, green, orange and pink, respectively.

### The age-dependent immune imprinting of anti-HA and anti-NA antibodies

With complete information on the HA and NA antigenic drift variants of A(H1N1) and A(H1N1)pdm09 viruses from the 1977 to the present time, we were able to compare the age-dependent anti-HA and anti-NA antibody profiles in the population. Subjects (N=130) aged 5-70 years (born between 1950 -2015) were recruited for a cross-sectional study with sera collected from April 2020 – January 2021, when there was minimal community transmission of seasonal influenza viruses due to increased public health and social measures against SARS-CoV-2 in Hong Kong (24). This unique sample set allowed us to profile the long-lasting anti-HA and anti-NA antibody responses without the interference of a recent influenza infection. WHO-recommended A(H1N1) (N=7) and A(H1N1)pdm09 (N=2) vaccine strains with distinct HA antigenicity were used to determine HI titers for all 130 participants.

We observed 96.2% (125/130) of the individuals developed ≥1:10 HI titer to a single or multiple HA antigens and 3.8% (5/130) did not show a detectable HI titer (<1:10) to any of the 9 HA antigens. By generating a heatmap using individually measured HI titers, we observed that 85.4% (111/130) of the participants developed ≥1:40 HI titers to at least one of the A(H1N1) or A(H1N1)pdm09 strains (Fig. 4a). Specifically, 17.7% (23/130) developed HI titer at ≥1:40 to a single antigen, and 67.7% (88/130) individuals developed cross-reactive HI antibodies to more than one antigen with HI titer ≥1:40.

**Figure 4.**
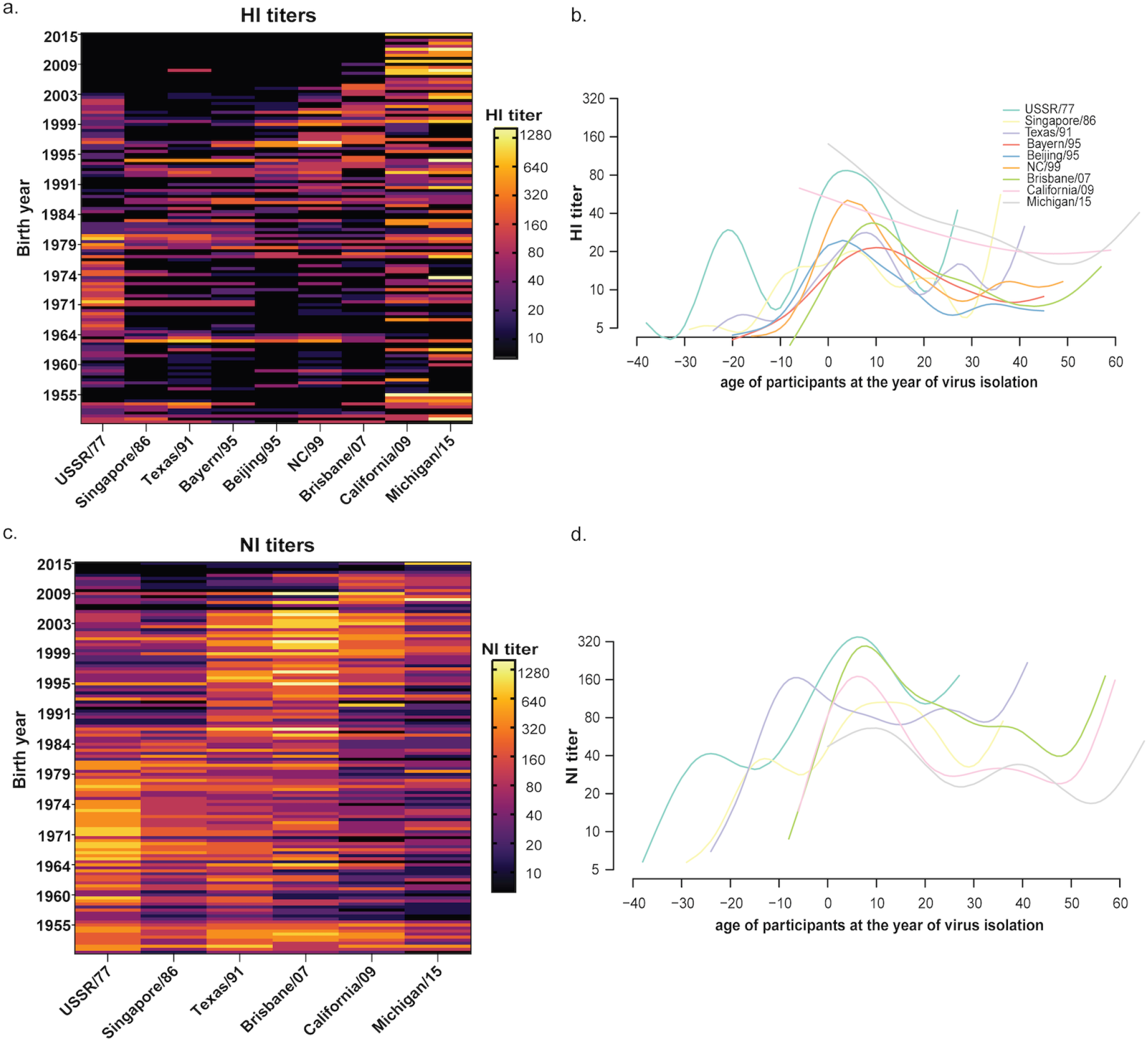
Age-dependent HI and NI antibody titers to antigenically distinct H1 HA and N1 NA of A(H1N1) and A(H1N1)pdm09 viruses from 1977 to 2015. **a**, A heatmap of 130 individuals’ HI titers against antigenically distinct A(H1N1) and A(H1N1)pdm09 viruses. The HI titers of each individual were represented in a row, sorted according to the birth year, with the youngest subjects shown on the top and the oldest subjects at the bottom. HI titers were shown in the log2 scale, with the light yellow shading indicating a higher HI titer and the black shading indicating a negative result (HI < 1:10). **b**, Generalised Additive Model (GAM) was fitted to the HI titers measured against A(H1N1) and A(H1N1)pdm09 viruses. The x-axis represents the age of each individual at the year of virus isolation of each A(H1N1) and A(H1N1)pdm09 viruses. The y-axis represents the HI titer. The fitted lines represent mean log2 HI titers against each A(H1N1) and A(H1N1)pdm09 viruses were color coded by the virus strain. Individuals showing HI titer <1:10 are arbitrarily assigned the titer 5. **c**, A heatmap generated using 130 individuals’ NI titers against antigenically distinct A(H1N1) and A(H1N1)pdm09 viruses. **d**, GAM was fitted to the NI titers measured against A(H1N1) and A(H1N1)pdm09 viruses. The x-axis represents the age of each individual at the year of virus isolation of each A(H1N1) and A(H1N1)pdm09 viruses. Individuals showing HI or NI titer <1:10 are arbitrarily assigned the titer 5.

To explore age-dependent imprinting against the HA protein of A(H1N1) and A(H1N1)pdm09 strains, generalised additive models (GAM) were fitted to distinct HI titers against the age of the participants at the year of strain isolation (Fig. 4b). The peak HI titers for most of the A(H1N1) strains were consistently observed in individuals aged 4-10 years old (∼7 years old) at the year of virus isolation. Interestingly, two HI peaks against USSR/77 were observed in individuals aged -21 years old (eg. born in 1998) or 4 years old (e.g. born in 1973) at the time when USSR/77 was isolated. No apparent peak was observed from the HI titers against the two A(H1N1)pdm09 viruses, California/09 and Michigan/15, suggesting the development of HI antibodies after primary A(H1N1)pdm09 infection was independent of birth year. These results confirm that early childhood exposure might lead to an age-dependent HI response against seasonal A(H1N1) viruses, while the HI response against A(H1N1)pdm09 viruses was less age-dependent.

Based on the knowledge of NA antigenic changes among A(H1N1) and A(H1N1)pdm09 strains, we further determine NI titers against these strains for all participants using ELLA. All 130 participants developed ≥1:10 NI titer to a single or multiple NA antigens, and most of the participants (95.4%) developed ≥1:40 NI titers to at least one of the antigenically distinct A(H1N1) and A(H1N1)pdm09 strains (Fig. 4c). Specifically, 6.2% (8/130) developed NI titer at ≥1:40 to a single NA antigen, and 90% (117/130) individuals developed cross-reactive antibodies to more than one NA antigen with NI titer ≥1:40. The percentage of individuals showing ≥1:40 cross-reactive NI titers was significantly higher than individuals showing ≥1:40 cross-reactive HI titers (p=0.000011, Chi-square test), suggesting that anti-NA antibodies may confer greater cross-reactivity than anti-HA antibodies.

By fitting NI titers against the age of the participants at the year of strain isolation, we observed that the peak NI titers for all the A(H1N1) strains were consistently observed from participants aged 6-12 years old (∼9 years old) at the year of virus isolation, with the exception of Texas/91 (Fig 4d). The peak NI titer for Texas/91 was observed among individuals -7 years old (eg. born in 1998) when the strain was isolated, suggesting the NA antigenicity has remained unchanged for an extended period of time, as reported previously (17). Interestingly, we observed age-dependent NI responses against A(H1N1)pdm09 viruses, with peak NI titers detected from participants age 6-10 years old at the time of virus isolation. This pattern was distinct from the age-independent HI response against A(H1N1)pdm09, as shown in Fig. 4b. The results suggest that the development of NI antibodies to A(H1N1) and A(H1N1)pdm09 viruses were age-dependent, with the primary exposure occurring within the first decade after birth leading to a long-lasting HI and NI imprint.

Overall, the serological assays have identified 125 of 130 individuals that developed detectable antibody titer (≥1:10) for both HI and NI analysis, with 50.7% (66/130) individuals showing the highest HI and NI titers to a single antigen, while 8.5% (11/130) showed both NI and HI titers for multiple antigens. Among these 77 individuals, only 42.9% (33/77) developed concordantly high HI and NI titers against the same virus strain. The correlation between the number of individuals showing the highest NI and HI titers to a single antigen and multiple antigens was calculated using a Chi-square test and found no significant association (p=0.225), irrespective of whether the individuals showed the highest titers to single or multiple antigens. These results support the discordant antigenic changes of influenza HA and NA proteins (16-18).

## DISCUSSION

We characterised the antigenic changes of N1 NAs of A(H1N1) from 1977 to 1991, which completed the NA antigenic characterisation of A(H1N1) and A(H1N1)pdm09 viruses circulating in humans since the 1950s, as the A(H1N1) virus re-emerged in 1977 was genetically related to A(H1N1) virus that circulated in the 1950s (25). From 1977 to 2020, changes in H1 antigenicity have been documented with a total of 13 WHO-recommended A(H1N1) (N=10) and A(H1N1)pdm09 (N=3) vaccine strains. During this time, only 6 N1 antigenic changes occurred in A(H1N1) (N=4) and A(H1N1)pdm09 (N=2) viruses based on current and previous studies (17, 18, 23), supporting the dissonant nature of antigenic drift of NA from HA (16-18, 23). By profiling the anti-HA and anti-NA antibody responses against this antigenically different A(H1N1) and A(H1N1)pdm09 viruses of 130 subjects born between 1950-2015, we observed age-dependent imprinting of both anti-HA and anti-NA antibody responses with the peak HI and NI antibodies generally detected from subjects 4-12 years old at the year of isolation of each antigenically distinct A(H1N1) strains. Interestingly, the HI response against the A(H1N1)pdm09 viruses was age-independent without apparent peak HI titer detected in any age group; however, the NI response against the A(H1N1)pdm09 was age-dependent with the peak NI titers detected from those 6-10 years old at the year of virus isolation. We also observed that 90% of the participants possessed ≥1:40 NI titers against more than one antigenically different NA protein, while only 67.7% of the participants possessed ≥1:40 HI titers against more than one antigenically different HA protein, suggesting that the antibodies generated to target the NA were more cross-reactive than those generated to target the HA receptor binding domain. Anti-NA antibodies have been established as a correlate of protection against influenza infection (4-7). As HA antigenic changes occur more frequently, anti-NA antibodies are expected to provide prolonged protection against HA drift variants if these strains continue to share NA antigenicity. Furthermore, the more cross-reactive nature of the anti-NA antibodies may provide partial protection against NA drift variants. Our results support the inclusion of NA antigen as a component of annual influenza vaccine preparation.

We identified that the NA of USSR/77, Singapore/86, and Texas/91 were antigenically distinct by two-way analysis, while there was a one-way antigenic change between the NA of USSR/77 and Chile/83. Kilbourne *et al*. previously reported antigenic stasis of NA in A(H1N1) vaccine strains between 1980-1983, but a one-way antigenic change was observed from the study (16). Similarly, one-way antigenic change was observed while characterising the N to K change at residue 386 from USSR/77 to Singapore/86, as changes in NI titers against USSR-N386K or Singapore-K386N were only observed with the use of heterologous ferret anti-sera. Previous studies by Sandbulte *et al*. and Gao *et al*. also identified that the antigenic drifts of N1 NAs were evident in one direction (18, 23). This raises the possibility that a one-way drift might not result in a complete loss of protection as the homologous anti-sera were able to tolerate a single amino acid change associated with antigenic drift. This also coincides with the observation that most of our study subjects possess ≥1:40 NI titers against more than one antigenically different NA protein. The broader breadth of NI responses can be supported by the conserved epitopes mapped by Chen *et al*. (N309 and N273) using mAbs on NAs across A(H1N1) and A(H1N1)pdm09 viruses from 1918 to 2017 (26). Similar findings were also observed by Wan *et al*. who recognised conserved mAb binding epitopes (group B mAbs identifying residues 273, 338, and 339) that were conserved across NAs of A(H1N1) and A(H1N1)pdm09 viruses from 1918 to 2009 (27).

The amino acid substitution N386K was identified to be responsible for the one-way antigenic drift from USSR/77 to Singapore/86. While N386 was preserved among A(H1N1) strains circulated from 1977 to 1984, A(H1N1) strains circulated from 1986 to 1988 harboured K386 and formed a separate cluster in the N1 phylogenetic tree. The subsequent K386D change found in A(H1N1) strains circulated from 1988 to 2008 may similarly result in NA antigenic drift from Singapore/86 to Texas/91. Interestingly, the N386K change was also reported to cause a one-way NA antigenic change in the A(H1N1)pdm09 viruses. The N386K change was reported to result in the loss of a potential glycosylation site and the NA antigenic drift between California/09 and Michigan/15 (18, 23). Pair-wise epistatic amino acid substitutions leading to changes in local net charges at the NA antigenic site have been implicated in maintaining viral fitness during evolution (28), and further research is needed to identify the role of epistatic changes in the proximity of residue 386. Taken together, these results showed N386K evolved independently from A(H1N1) and A(H1N1)pdm009 viruses and suggested convergent evolution in N1 NA of human influenza viruses.

Age-dependent imprinting of anti-HA and anti-NA antibody responses was observed from 130 subjects born between 1950–2015, with peak HI and NI antibodies generally detected from subjects at 4-12 years old when each of the antigenically distinct A(H1N1) strains was isolated. This finding is in accordance with previous studies showing age-dependent HI responses. Lessler *et al*. estimated that virus neutralisation titers measured against a panel of A(H3N2) viruses were the highest in subjects aged ∼7 years at the time of strain isolation and declined smoothly thereafter across all strains (29). A similar study by Yang *et al*. estimated the highest HI titers against a panel of A(H3N2) viruses at a median age of 4.3 years (IQR, 2.0-6.9 years) during strain isolation (30). Overall, the results suggest that primary exposure to seasonal influenza viruses occurred within the first decade of life, which is in agreement with the higher infection attack rates of seasonal influenza viruses in children than in adults (31, 32). Interestingly, the HI response against the A(H1N1)pdm09 viruses did not show an apparent age-dependent pattern. A higher infection attack rate of A(H1N1)pdm09 virus was similarly reported in children than in adults during the first year of the pandemic (33, 34). However, an increasing proportion of adults were infected by the A(H1N1)pdm09 virus in subsequent years (35), which may have weakened the age-dependent HI response against the A(H1N1)pdm09 virus. In comparison, the NI responses against the A(H1N1)pdm09 viruses were age-dependent, with peak NI titers detected from those at 6-12 years old at the time of virus isolation. It is unclear if anti-NA antibodies raised against A(H1N1) viruses conferred cross-protection against A(H1N1)pdm09 virus in adults, while the age-dependent anti-NA antibodies against A(H1N1)pdm09 were developed among those who experience A(H1N1)pdm09 as the first influenza infection in life. Our study is limited by the vaccination history and recent infection history of the study subjects. However, sera were collected during 2020-2021 when minimal influenza activity in the community may have reduced the interference of recent influenza infection on HI and NI responses.

HI and NI antibody responses continue to be shaped by repeated infection and vaccination with antigenically similar or distinct strains during our lifetime. Our study confirmed the age-dependent HI and NI responses and identified the more broadly cross-reactive nature of anti-NA antibodies than the anti-HA antibodies. Future longitudinal studies where individuals are followed up since birth could provide better insights into protection conferred by the anti-HA and anti-NA antibodies. Our study supports the inclusion of NA protein into annual influenza vaccine preparation and emphasises the importance of routinely monitoring of NA antigenic drifts in parallel with HA evolution.

## MATERIALS & METHOD

### Cell culture

Madin-Darby canine kidney (MDCK) cells and Human Embryonic Kidney (HEK) 293T cells purchased from American Type Cell Culture (ATCC) were grown in Minimum Essential Media (MEM) and Opti-MEM, respectively (Gibco). The MEM used for cell culture was supplemented with 10% fetal bovine serum (Gibco), Penicillin -Streptomycin (Gibco), Vitamins (Sigma-Aldrich) and HEPES (Gibco). The infection media (MEM, Gibco) used for viral infection was supplemented with 0.3% Bovine Serum Albumin (Sigma-Aldrich), Penicillin -Streptomycin (Gibco), Vitamins (Sigma-Aldrich) and HEPES (Gibco).

### Viruses

Recombinant H6N1 viruses were generated for serological analysis using plasmid-based reverse genetics, as previously described (36). The NA genes of A(H1N1) vaccine strains for the period 1977-1991; A/USSR/90/77, A/Chile/01/83, A/Singapore/06/86 and A/Texas/36/91, A/Brisbane/59/07, A/California/04/09-pdm09 and A/Michigan/45/15-pdm09 were RT-PCR amplified and cloned into the pHW2000 vector. The HA gene of A/Teal/Hong Kong/W312/97 (H6N1) has been cloned into pHW2000 as described previously (37). The HA and NA plasmids were co-transfected with pHW181, pHW182, pHW183, pHW185, pHW187 and pHW188 derived from A/Puerto Rico/8/34 (PR8) in 293T cells using TransIT (Mirus) and Opti-MEM (Gibco) to generate recombinant H6N1 viruses. Transfection supernatants were harvested and passaged in MDCK cells at an MOI of 0.001 PFU/cell to prepare stock viruses. The HA and NA genes of all stock viruses were sequence verified using sanger sequencing prior to use.

The egg-passaged A(H1N1) viruses, A/USSR/90/77, A/Singapore/06/86, A/Texas/36/91, A/Bayer/272/95, A/Beijing/07/95, A/New Caledonia/20/99, A/Brisbane/59/07, A/California/04/09-pdm09 & A/Michigan/45/15-pdm09 were used in the HI assay were kindly supplied by the Centre for Disease Control and Prevention (CDC), Atlanta, GA and Francis Crick Institute, Midland, London. The viruses were propagated in 9-11 day old specific pathogen-free (SPF) embryonated chicken eggs. The allantoic cavity of each egg was injected with 100 μl of virus diluted in PBS, supplemented with Penicillin-Streptomycin (Gibco) and Gentamicin (Gibco). The site of virus injection was sealed, and the eggs were incubated at 37 °C for 48 hours. The eggs were then transferred to 4 °C to be kept overnight before harvesting the virus.

### Ferret anti-sera

Ferret anti-sera raised against wild-type A(H1N1) viruses; A/USSR/90/77, A/Hong Kong/117/77, A/Brazil/11/78, A/Lackland/3/78, A/Ostrava/12/80, A/Dakar/4363/81, A/Finland/10/81, A/Firenze/13/83, A/Chile/01/83, A/Victoria/4/84, A/Singapore/06/86, A/Taiwan/1/86, A/Sichuan/4/88 and A/Texas/36/91 viruses were generously provided by Francis Crick Institute (Midland, London) and by CDC (Atlanta, GA).

### Study group and serum samples

Human serum for the cross-sectional study was collected from blood donors at 18-70 years of age (N=110) from April-August 2020 (IRB number UW-132). The pediatric samples were collected for a study on SARS-CoV-2 infection in patients aged 5-17 years (N=20) from May 2020 – February 2021 (UW 21-093) during the symptom onset of day 0 to 7 months as a part of a previous study. A total of 130 individuals aged 5-70 were included in the serology study.

### ELLA to measure the Neuraminidase Inhibiting antibody (NI) Titers

NI antibody titers were measured as previously described (20). First, the dilution of the virus that resulted in a 90–95% maximum signal was elected for use in serology. The sera were heat treated (56 °C for 45 min), 2-fold serially diluted in PBS–BSA and added to duplicate wells on a fetuin (Sigma-Aldrich) coated plate. An equal volume of the selected virus dilution was added to all serum-containing wells in addition to wells containing diluent without serum that served as a virus-only control. The plates were incubated for 16–18 hrs at 37 °C, then washed with PBS– 0.05% Tween 20 (PBS-T) before adding 100 μl/well peanut agglutinin conjugated to horse-radish peroxidase (PNA-HRPO, Sigma-Aldrich). Plates were incubated at room temperature for 2 hrs and washed with PBST before adding o-phenylenediamine dihydrochloride (OPD, Sigma-Aldrich) to the plate. A 100 μl of the OPD substrate was added to each well on all plates and incubated in the dark, and the colour reaction was stopped by adding 100 μl/well of 1N sulfuric acid. The plates were read at 490 nm for 0.1 s using a Microplate Fluorimeter (FLUOstar OPTIMA F, BMG LABTECH).

### HI assay to measure the anti-HA antibody titers

Human sera were treated with Receptor Destroying Enzyme (RDE) II (Accurate, #YCC340-122) overnight and were heat inactivated for 30 min at 56 °C. The heat-inactivated sera were serially 2-fold diluted and incubated with A(H1N1) and A(H1N1)pdm09 viruses diluted to 8 HA/50μl for 30 min at room temperature. 0.5% Turkey RBCs were added to the mixture and incubated for 30 min. The highest sera dilution that inhibited hemagglutination was recorded as the HI titer.

### Antigenic cartography

Antigenic cartography of end-point ELLA two-way NI titers (Table 1) generated using recombinant influenza A(H6N1) viruses and polyclonal A(H1N1)-raised ferret antisera was optimized over 1000 iterations into a two-dimensional (2D) antigenic landscape with the ACMACS-API software suite (version acmacs-c2-20161026-0717 and i19 build host) (38). NI titer values of 5 were assigned for all results below the ELLA limit-of-detection (NI titer < 10). Modelled NI trends were rendered into 2D X/Y-mapping coordinates and annotated in Tableau Desktop (version 2022.3.0). All applied code sets are available by the authors upon request.

### Genetic analysis

The phylogenetic tree was constructed using the full coding region of the NA gene of 89 selected human A(H1N1) strains from 1977 to 2008 obtained from Global Initiative on Sharing All Influenza Data (GISAID). NA nucleotide sequences were aligned, and the maximum-likelihood tree (bootstrap 500 replicates) was constructed using MEGA (version 11.0.10). The constructed tree was visualised with the Geneious Prime® 2023.0.1

### Site-directed mutagenesis

Single amino acid changes, S247N, K369R, N386K and K434N, were introduced into the plasmid encoding USSR/77-NA gene, and the amino acid change K386N was introduced into the plasmid encoding Singapore/86-NA gene using the QuikChange II site-directed mutagenesis kit (Agilent Technologies).

### Statistical analysis

RStudio (version 1.3.1093) was used to generate GAM fitted to log2 HI and NI titer against the age of the individuals at the time of virus isolation using the mgcv package (version 1.8-33).

## ACKNOWLEDGEMENTS

This study was supported by RGC Theme-based Research Schemes, Hong Kong SAR, China (T11-705/14N and T11-712/19-N). The authors would like to thank Dr. Ranawaka APM Perera from HKU for helpful discussions, Samuel S. Shepard from Centers for Disease Control and Prevention and Reina Chau from General Dynamics Information Technologies, Inc. for their contributions in operationalizing ACMACS for portable and high-throughput usage.

